# A genome-wide association study finds genetic variants associated with neck or shoulder pain in UK Biobank

**DOI:** 10.1101/2020.01.20.913228

**Authors:** Weihua Meng, Brian W Chan, Cameron Harris, Maxim B Freidin, Harry L Hebert, Mark J Adams, Archie Campbell, Caroline Hayward, Hua Zheng, Xianwei Zhang, Lesley A Colvin, Tim G Hales, Colin NA Palmer, Frances MK Williams, Andrew McIntosh, Blair H Smith

## Abstract

**Background:** Common types of musculoskeletal conditions include pain in the neck and shoulder areas. This study seeks to identify the genetic variants associated with neck or shoulder pain based on a genome-wide association approach using 203,309 subjects from the UK Biobank cohort and look for replication evidence from the Generation Scotland: Scottish Family Health Study (GS:SFHS) and TwinsUK.

**Methods:** Cases in the UK Biobank were determined by a question which asked the participants if they had experienced pain in the neck or shoulder in the previous month influencing daily activities. Controls were the UK Biobank participants who reported no pain anywhere in the last month. A genome-wide association study was performed adjusting for age, sex, BMI and 9 population principal components. Significant and independent genetic variants were then sent to GS:SFHS and TwinsUK for replication.

**Results:** We identified 3 genetic loci that were associated with neck or shoulder pain in the UK Biobank samples. The most significant locus was in an intergenic region in chromosome 17, rs12453010, having *P* = 1.66 × 10^-11^. The second most significant locus was located in the *FOXP2* gene in chromosome 7 with *P* = 2.38 × 10^-10^ for rs34291892. The third locus was located in the *LINC01572* gene in chromosome 16 with *P* = 4.50 × 10^-8^ for rs62053992. In the replication stage, among 4 significant and independent genetic variants, rs2049604 in the *FOXP2* gene and rs62053992 in the *LINC01572* gene were weakly replicated in GS:SFHS (*P =* 0.0240 and *P* = 0.0202, respectively). None of the single nucleotide polymorphisms (SNPs) were replicated in the TwinsUK cohort (*P* > 0.05).

**Conclusions:** We have identified 3 loci associated with neck or shoulder pain in the UK Biobank cohort, two of which were weakly supported in a replication cohort. Further evidence is needed to confirm their roles in neck or shoulder pain.

**Significance:** This is the first genome-wide association study on neck or shoulder pain. We have identified 3 genetic loci (an intergenic region in chromosome 17, the *FOXP2* gene in chromosome 7, and the *LINC01572* gene in chromosome 16) that are associated with neck or shoulder pain using the UK Biobank cohort, among which the *FOXP2* gene and the *LINC01572* gene were weakly replicated by the Generation Scotland: Scottish Family Health Study (*P* < 0.05). The SNP heritability was 0.11, indicating neck or shoulder pain is a heritable trait. The tissue expression analysis suggested that neck or shoulder pain was related to multiple brain tissues, indicating the involvement of neuron function. The results will inform further research in the characterisation of the mechanisms of neck or shoulder pain.

## INTRODUCTION

Musculoskeletal pain in the neck and shoulder areas is a major health problem for adults of working age as well as for elderly populations (1). Neck and shoulder pain are prevalent forms of self-reported musculoskeletal pain (2). The aetiologies of neck and shoulder pain may be complicated since both regional lesions and systemic disorders outside the cervicobrachial area may cause pain at that location (3, 4). In addition, lesions in the neck can lead to pain in the shoulder and vice versa (5). Many people also have difficulty in describing and differentiating pain in these areas accurately. For these reasons, neck or shoulder pain is often discussed as a single entity (6).

Epidemiological studies have suggested that the prevalence of neck pain is 6-22% and 7-27% for shoulder pain (7–9). The Global Burden of Disease Study 2010 found that of the 291 conditions studied, neck or shoulder pain as a single entity ranked 21st in overall burden on society, and 4th in terms of overall disability (8). The updated Global Burden of Disease Study 2016 also indicated that neck pain was a top five cause of years lived with disability (YLD) in high-income, and high-middle-income countries.(10) Risk factors associated with neck or shoulder pain conform to the biopsychosocial model; specifically, they include older age, being female, high body mass index (BMI), previous injury, strenuous occupation and diabetes mellitus (3,11–14). Although mechanical exposure is associated with increased risk of pain in the neck and shoulder, this explains only part of these complaints (15). Because of the biopsychosocial factors involved, treating neck or shoulder pain successfully is a challenge. In a study of neck, shoulder and arm pain, only 25% of the patients made a complete recovery after 6 months (16). Estimated rates of remission 1 year after neck or shoulder pain onset were between 33-55% (17–20).

Genetic studies have identified genes associated with neck or shoulder pain. Twin studies have shown that there is a genetic role in neck pain (14), though in keeping with many traits the genetic component becomes smaller with age (21, 22). Nonetheless, in adolescents as much as 68% of variance in neck pain liability could be attributed to genetic factors (23). So far there has been no genome-wide association study (GWAS) published on neck or shoulder pain.

This study seeks to identify the genetic variants associated with neck or shoulder pain based on a GWAS approach in a cohort of 203,309 subjects from the UK Biobank cohort and to test significant results for replication in the Generation Scotland: Scottish Family Health Study (GS:SFHS) and TwinsUK. Similar approaches have been used to examine back pain, knee pain, and headaches in the UK Biobank cohort (24–26).

## METHODS

### Participants and the genetic information of cohorts

Discovery cohort - The UK Biobank (https://www.ukbiobank.ac.uk/) is a project facilitating research into health and disease and involves over 500,000 participants aged between 40-69 years old at recruitment. Participants completed a detailed questionnaire which examined lifestyle, demographic factors, and clinical history. Participants also underwent clinical measures and baseline body measurements such as height and weight. Biological samples including urine, saliva, and blood were also provided. The National Research Ethics Service granted ethical approval to the UK Biobank (reference 11/NW/0382). The genetic information of 500,000 participants was released to approved researchers in March 2018. The corresponding author of this paper was granted access to the genetic information under UK Biobank application number 4844. Detailed quality control information pertaining to these genotypes was described by Bycroft et al (27).

Replication cohort 1 - Generation Scotland: Scottish Family Health Study (GS:SFHS) is a multi-institutional, family-based cohort involving over 20,000 volunteer participants aged between 18-98 years old at recruitment, who provided blood samples from which DNA was extracted. Similarly, participants also completed questionnaires to provide detailed phenotypic and sociodemographic information. Clinical and biochemical measurements were also collected for the purpose of research. Permission was obtained for linkage of research data to routine health data in the form of electronic health records (28, 29). Ethical approval for GS:SFHS was obtained from the Tayside Committee on Medical Research Ethics (on behalf of the National Health Service) with reference Number 05/S1401/89. The genetic information relating 20,000 participants was released to the corresponding author of this paper in March 2018 for pain-related research. Detailed quality control information pertaining to these genotypes was described by Hall et al (30). Replication cohort 2 - The TwinsUK cohort is a UK nationwide registry of volunteer same sex twins. It has recruited 14,274 registered twins aged between 16 and 98 years. Collection of data and biological materials commenced in 1992 and is ongoing. During study participation, participants regularly complete health and lifestyle questionnaires and visit collaborating clinics and hospitals for clinical evaluation. Ethical approval was provided by the Research Ethics Committee at Guy’s and St. Thomas’ NHS Foundation Trust. TwinsUK has the genetic information relating to 6,921 participants. Detailed quality control information pertaining to the genetic information was described by Moayyeri et al (31).

### Phenotypic definitions on neck or shoulder pain

Discovery cohort - UK Biobank: Participants were offered a pain-related questionnaire, which included the question: ‘in the last month have you experienced any of the following that interfered with your usual activities?’. The options were: 1. Headache; 2. Facial pain; 3. Neck or shoulder pain; 4. Back pain; 5. Stomach or abdominal pain; 6. Hip pain; 7. Knee pain; 8. Pain all over the body; 9. None of the above; 10. Prefer not to say. Participants could select more than one option. (UK Biobank Questionnaire field ID: 6159).

In this study, cases were defined as participants who reported having activity limiting pain in the neck or shoulder in the past month (option 3), regardless of whether they reported pain in other regions. The controls were defined as participants who chose the ‘None of the above’ option.

Replication cohort 1 – GS:SFHS: Participants were first asked ‘’Are you currently troubled by pain or discomfort, either all the time or on and off?’’. If yes was selected, then the participants were asked ‘’Have you had this pain or discomfort for more than 3 months?’’. If yes was selected once again, then they were asked ‘’Where is this pain or discomfort?’’ with options of ‘Back pain’, ‘Neck or shoulder pain’, ‘Headache, facial or dental pain’, ‘Stomach ache or abdominal pain’, ‘Pain in your arms, hands, hips, legs or feet’, ‘Chest pain’, and ‘Other pain’. If a participant selected the ‘Neck or shoulder pain’, then he/she was defined as a case. All other subjects were defined as controls.

Replication cohort 2: TwinsUK: participants were asked ‘In the past three months, have you had pain in your neck or shoulders?’ Those who answered ‘Yes’ were defined as cases. Those who answered ‘No’ were defined as controls. Those with missing answers were not included in the study.

### Statistical Analysis

Discovery cohort – UK Biobank: BGENIE (https://jmarchini.org/bgenie/) was used as the main GWAS software. Single nucleotide polymorphisms (SNPs) with imputation INFO scores < 0.1, minor allele frequency (MAF) < 0.5% were removed, as well as SNPs that failed Hardy-Weinberg tests *P* < 1.0 × 10^-6^.

BGENIE was used to perform association studies using linear association tests, adjusting for age, sex, BMI, 9 population principle components, genotyping arrays, and assessment centres. Chi-square testing was used to compare gender difference between cases and controls. T-tests were used to compare age and BMI between case and control groups using IBM SPSS 22 (IBM Corporation, New York). As is standard in GWAS, SNP associations were considered significant when *P* < 5.0 × 10^-8^. Genome-wide Complex Trait Analysis (GCTA) software was used to calculate SNP-based or narrow-sense heritability (32). Significant and independent SNPs from the GWAS results of the discovery cohort were then sent to replication cohorts for replication. These significant and independent SNPs were defined by FUMA with r^2^ (linkage disequilibrium score) < 0.6 with any other significant SNPs.

Replication cohort 1 – GS:SFHS: GCTA fastGWA1.92.4 was the main software used for replication. (https://cnsgenomics.com/software/gcta/#fastGWA). SNPs with INFO scores < 0.3, or MAF < 1% were removed, as well as SNPs that failed Hardy-Weinberg tests *P* < 1.0 × 10^-6^. FastGWA was used to perform association studies using a mixed-effects linear model adjusting for age, sex, BMI, and 9 population principle components. Relatedness was adjusted for via a genetic kinship matrix. Replication cohort 2 – TwinsUK: GEMMA v 0.98.1 was used for replication (https://github.com/genetics-statistics/GEMMA). A mixed-effects linear model adjusting for age, sex, BMI, and relatedness via a genetic kinship matrix was used. Meta-analysis of the significant and independent SNPs combining the UK Biobank, GS:SFHS and TwinsUK was performed using GWAMA 2.2.2 (https://genomics.ut.ee/en/tools/gwama).

Post-GWAS analysis: This study used FUMA as a main annotation tool for viewing and annotating GWAS results (33). It applied SNP functional annotations and generated a corresponding GWAS Manhattan plot.

MAGMA v1.06 (integrated in FUMA) was used to perform gene-based association analysis and gene-set analysis, both of which were generated from GWAS summary statistics (34). For gene-based association analysis, all SNPs located in protein coding genes are mapped to one of 19,123 protein coding genes. The default SNP-wise model (mean) was applied. We tested the joint association of all SNPs in the gene with the phenotype by aggregating the SNP summary statistics to the level of whole genes. In gene-set analysis, individual genes were aggregated to groups of genes sharing certain biological, functional or other characteristics. This aims to elucidate the involvement of specific biological pathways or cellular functions in the genetic aetiology of a phenotype. GTEx (also integrated in FUMA, https://www.gtexportal.org/home/) provided the results of tissue expression analysis.

Genetic correlation analysis was also performed to identify genetic correlation between neck or shoulder pain and 234 complex traits based on the online tool LD hub v1.9.0 (http://ldsc.broadinstitute.org/ldhub/). Any *P* value less than 2.1 × 10^-4^ (0.05/234) was considered statistically significant by Bonferroni adjusted testing.

## RESULTS

### GWAS Results

In the UK Biobank, 775,252 responses to all options were received for the specific pain question. Of the 501,708 participants in the study, 123,061 participants reported having experienced activity limiting pain in the neck or shoulder in the previous month. 213,408 participants chose the ‘None of the above’ option which meant they did not have activity limiting pain anywhere in the previous month. To create a homogeneous dataset, we first removed samples according to their ancestry information. In addition, those who were related to one or more others in the cohort (a cut-off value of 0.044 in the generation of the genetic relationship matrix) and those who failed quality control were also removed. The final number of those included in the case group after the above exclusions was 53,994 (28,093 males, 25,901 females). 149,312 (71,480 males, 77,832 females) individuals were included in the control group. After SNP quality control, there were 9,304,965 SNPs available for GWAS analysis. Clinical characteristics of the case and control groups were compiled (Table 1). Age, sex, and BMI were all found to be significantly different (*P* < 0.001) between cases and controls. Three genetic loci including 4 significant and independent SNPs reached a GWAS significance of *P* < 5×10^-8^ (Figure 1, Table 2). The most significant locus was located in an intergenic region in chromosome 17. The SNP from this location of highest significance was rs12453010 (*P* = 1.66 × 10^-10^). The second locus was found in the *FOXP2* gene located in chromosome 7, and the most significant SNP from this locus was rs34291892 (*P* = 2.38 × 10^-10^). The third locus was the *LINC01572* gene located in chromosome 16, and the most significant SNP in this locus was rs62053992 (*P* = 4.50 × 10^-10^). The SNP heritability (liability scale) from GCTA was 0.11±0.017.

**Figure 1:**
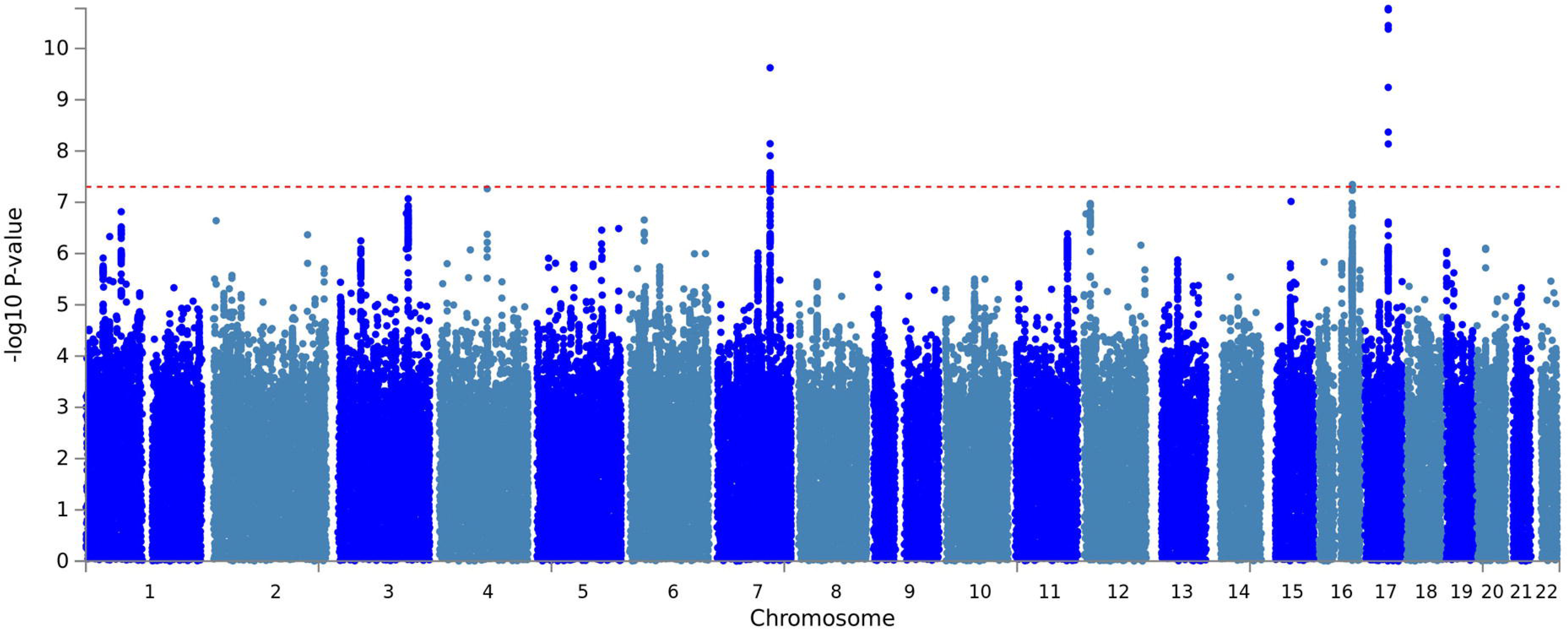
The Manhattan plot of the GWAS on neck or shoulder pain using the UK Biobank cohort (N=203,309). The dashed red line indicates the cut-off *P* value of 5 × 10^−8^.

**Table 1.** Clinical characteristics of neck or shoulder pain cases and controls in the UK Biobank

| Covariates | UK Biobank |  | *P* |
| --- | --- | --- | --- |
|  | Cases | Controls |  |
| Sex (male:female) | 28093 : 25901 | 71,480 : 77,832 | <0.001 |
| Age (years) | 57.7 (7.82) | 56.9 (7.97) | <0.001 |
| BMI (kg/m<sup>2</sup>) | 27.8 (4.85) | 26.7 (4.30) | <0.001 |

BMI: body mass index
A chi-square test was used to test the difference of gender frequency between cases and controls and an independent t test was used for other covariates.
Continuous covariates were presented as mean (standard deviation).

**Table 2.**
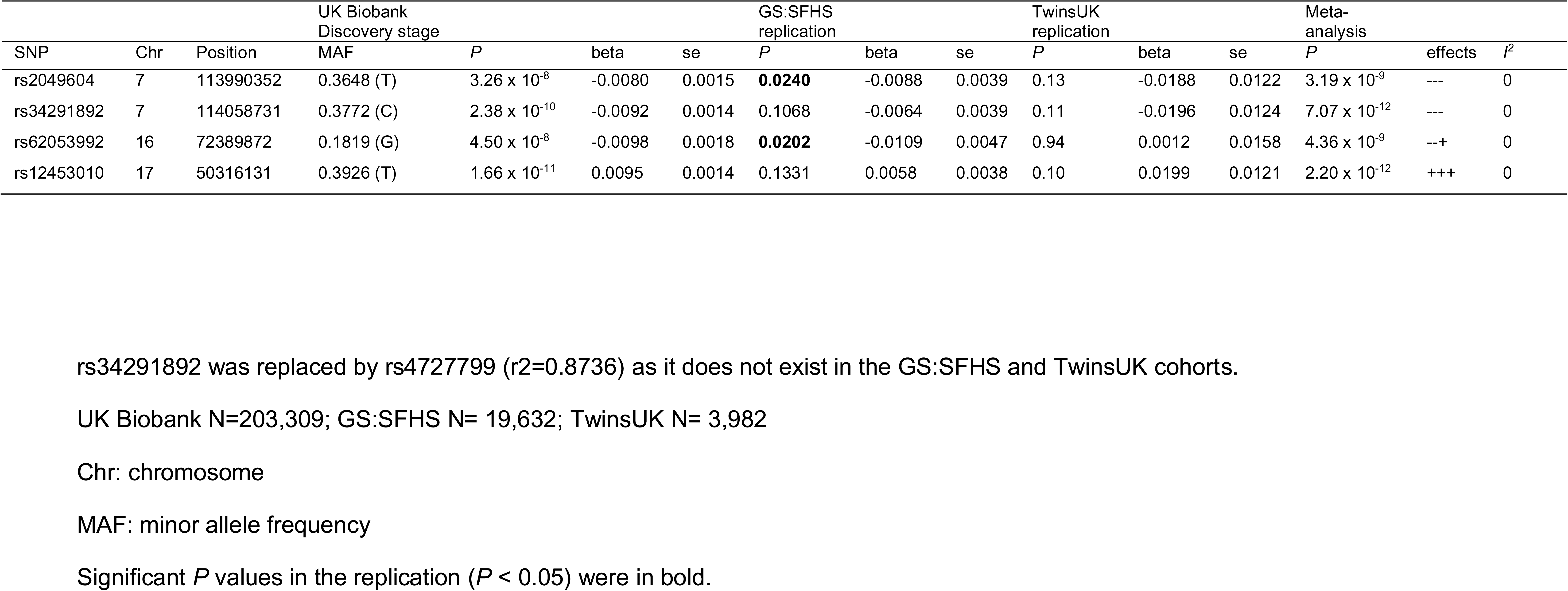
The summary statistics of the 4 significant and independent SNPs in the UK Biobank and the replication and meta-analysis results using the GS:SFHS and TwinsUK cohorts

These 4 significant and independent SNPs were tested for replication in GS:SFHS and TwinsUK. Among 20,032 subjects in the GS:SFHS subjects, 19,632 had complete relevant information. The whole-genome fastGWA results did not find any SNPs with GWAS significance associated with neck or shoulder pain. (Supplementary Figure S1). Among the 4 SNPs from the discovery cohort, rs2049604 in the *FOXP2* gene and rs62053992 in the *LINC01572* gene were replicated weakly (*P* = 0.0240 and 0.0202, respectively) (Table 2). Of the 6,921 individuals in TwinsUK with genetic information, 3,982 of them had valid relevant information on phenotypes and covariates. None of the 4 SNPs were replicated in the TwinsUK cohort (*P* > 0.05).

Meta-analysis of the 4 significant and independent hits, combining UK Biobank, GS:SFHS and Twins UK found that the significance of their associations was increased (Table 2).

### FUMA Analysis

In gene-based association analysis by MAGMA, 26 genes were found to be associated with neck or shoulder pain, all of which are represented in Supplementary Table S1. The most significant gene was *FOXP2* (*P* = 1.62 × 10^−11^), which is located in chromosome 7.

In gene-set analysis conducted by MAGMA, 10,651 gene-sets were analysed using the default competitive test model. None of these gene-sets met genome-wide significance *(P* < 4.7 × 10^−6^ (0.05/10651)).

Tissue expression analysis was conducted by GTEx, and the relationship between tissue specific gene expression and genetic associations was tested by using the average gene expression in each tissue type as a covariate. Two analyses were carried out, one investigating 30 general tissue types (Figure 2) and the other looking at 53 specific tissue types (Figure 3). Tissue expression analysis in 30 tissue types found expression in brain tissue to be the most significant (*P* = 9.53 × 10^−5^). Only expression in brain and pituitary tissue reached significant values of *P* < 1.67 × 10^−3^ (0.05/30). Tissue expression analysis of 53 specific tissue types by GTEx found expression in the nucleus accumbens of the basal ganglia to be the most significant (*P* = 3.55 × 10^−5^). In addition, the top 6 significant associations were all from brain tissues.

**Figure 2:**
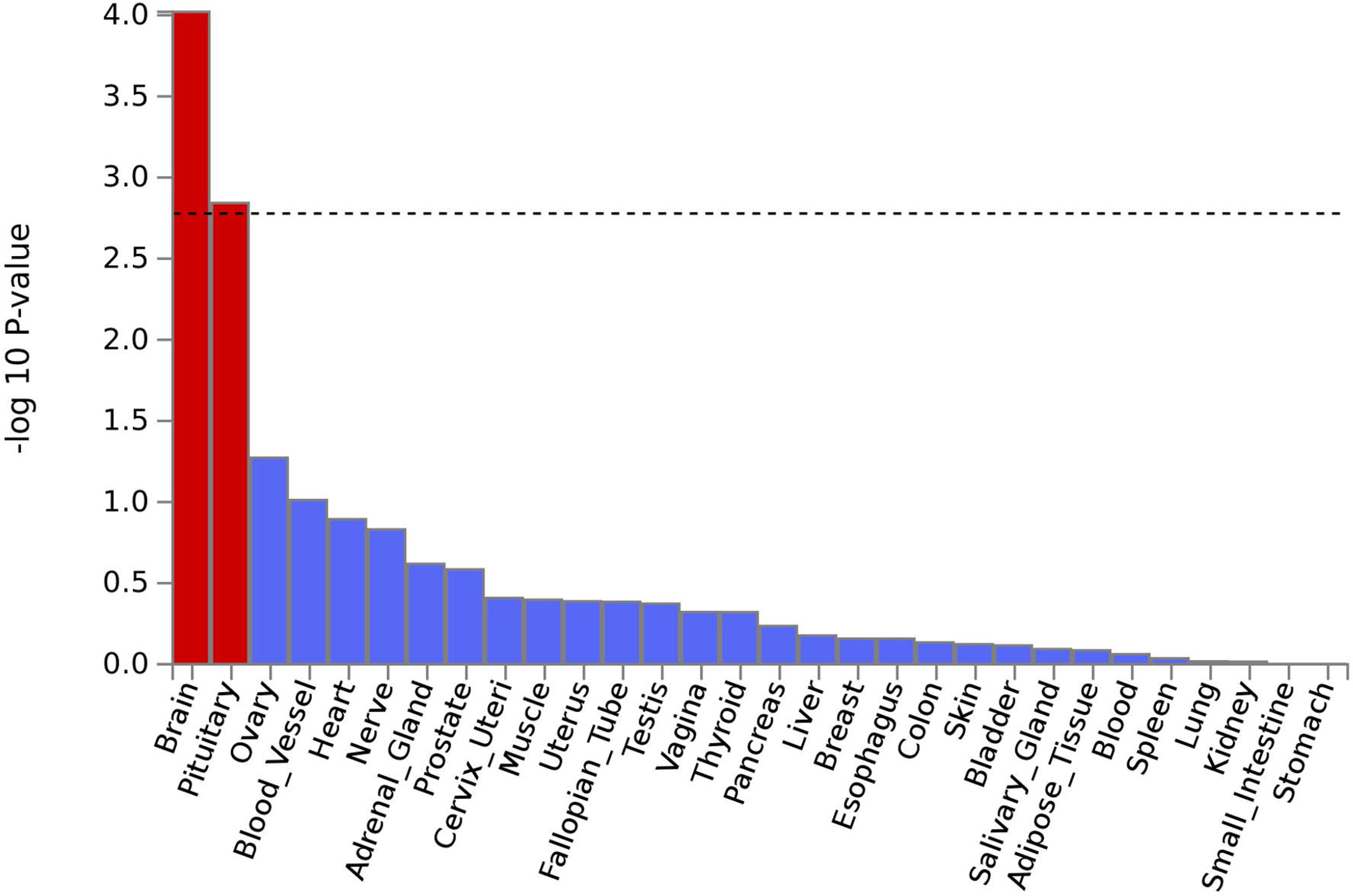
Tissue expression results on 30 specific tissue types by GTEx in the FUMA. The dashed line shows the cut-off *P* value for significance with Bonferroni adjustment for multiple hypothesis testing.

**Figure 3:**
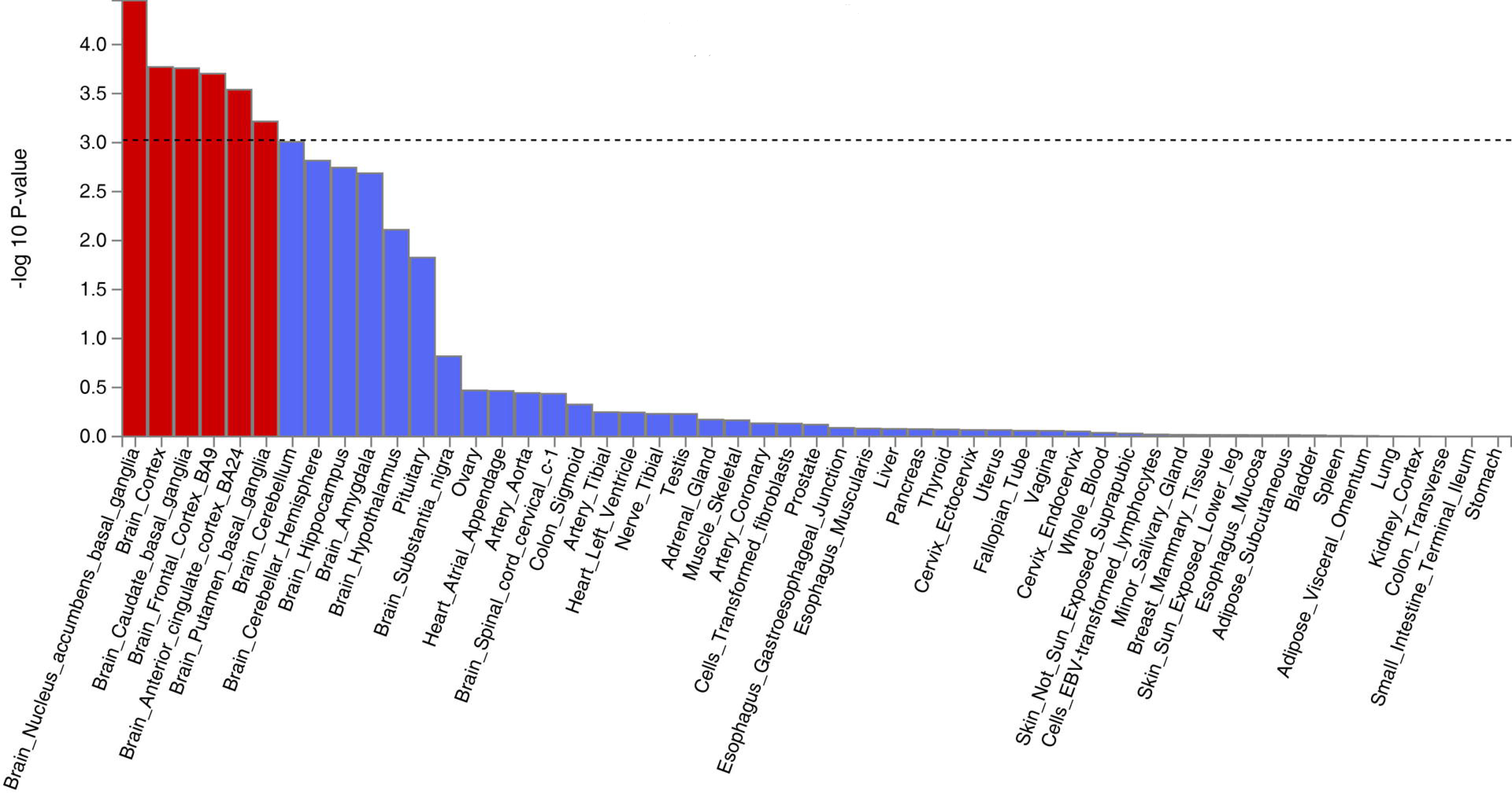
Tissue expression results on 53 specific tissue types by GTEx in the FUMA. The dashed line shows the cut-off *P* value for significance with Bonferroni adjustment for multiple hypothesis testing.

The genetic correlation analysis using the LD hub (v1.9.0) showed that symptoms of depression had the largest significant and positive genetic correlation with neck or shoulder pain (rg = 0.5522, *P* = 3.41 × 10^−30^), followed by insomnia (rg = 0.5377, *P* = 1.21 × 10^−21^). It was also found that the age at which they have their first child had the most significant and negative genetic correlations with neck or shoulder pain (rg = -0.4812, *P* = 8.95 × 10^−37^), followed by college completion (rg = -0.4706, *P* = 7.26 × 10^−26^). All the phenotypes with significant genetic correlation with neck or shoulder pain are shown in Table 3.

**Table 3.**
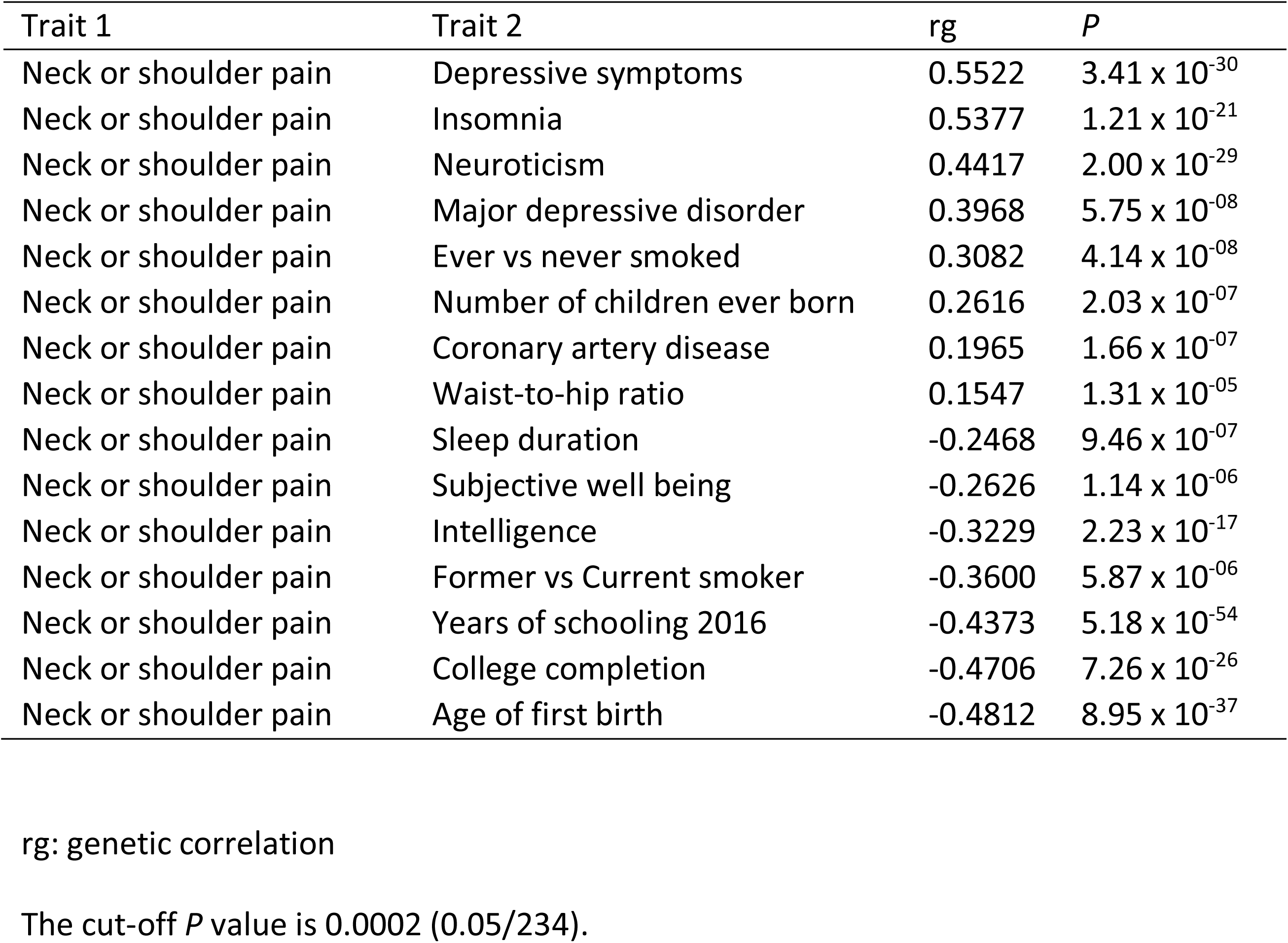
The significant genetic correlation results by the LD hub between neck or shoulder pain with other phenotypes

## DISCUSSION

We have performed a GWAS on neck or shoulder pain using the UK Biobank resource and found 3 loci that have reached genome-wide significance (*P* <0.05 × 10^−8^). They are the *FOXP2* gene in chromosome 7, the *LINC01582* gene in chromosome 16 and an intergenic area in chromosome 17. In the replication stage, the *FOXP2* and *LINC01582* loci were weakly supported by the GS:SFHS cohort, but not by TwinsUK.

*FOXP2* belongs to the forkhead-box transcription factor family and encodes a 715 amino acid long transcription factor (35). It may have 300-400 transcription targets, and has a forkhead/winged helix binding domain with 2 polyglutamine tracts adjacent to each other due to a mixture of CAG and CAA repeats (36, 37). *FOXP2* is a gene shown to be vital in the neural mechanisms underpinning the development of speech and language. A previous study described a family with developmental verbal dyspraxia, where affected individuals had a loss of function mutation in the *FOXP2* gene (38). Affected individuals in the family had difficulty with the selection and sequencing of fine orofacial muscular movements needed to articulate words, as well as deficits in language and grammatical skills. *FOXP2* also plays a role in regulating ‘hub’ genes *Dlx5* and *Syt4* in animal models, which are important for brain development and function (39–41). The mutations in the *FOXP2* gene were also associated with decreased grey matter in the cerebellum (42).

Notably, *FOXP2* is expressed in several regions of the brain, namely the basal ganglia, locus coeruleus, parabrachial nucleus, and thalamus. All of these regions have been previously implicated in the modulation of pain (43–46), and also concur with the tissue expression analysis (Figure 2, Figure 3) which suggests that the central nervous system modulates neck or shoulder pain. A recent GWAS study suggested that *FOXP2* is a candidate gene for multisite chronic pain (47). Further studies into the location and function of the transcription targets of *FOXP2* would also provide valuable insight.

*The LINC01572* gene in chromosome 16 was also replicated in this study, with rs62053992 having the *P* value at 4.5×10^−8^ in the discovery cohort and at 0.02 in the GS:SFHS. The gene is 384kb long, and was recently suggested to be related with polycystic ovary syndrome (48). However, studies of this gene have been very limited. Further replication evidence should be sought to confirm its role in neck or shoulder pain. Although the locus in chromosome 17 was not replicated, it was the most significantly associated in the discovery cohort with rs12453010 having the lowest *P* value of 1.66 × 10^−11^. Analysis of this locus is also difficult as it is a gene desert area. The nearest neighbouring gene is *CA10,* which encodes a protein belonging to the carbonic anhydrase family and is responsible for catalysing the hydration of carbon dioxide. It is also thought to contribute to central nervous system development particularly in the brain (49).

The results of the meta-analysis of the 4 significant and independent SNPs combining 3 cohorts suggested that the loci identified by the UK Biobank cohort were supported by the GS:SFHS and TwinsUK. It is likely that the sample size in the TwinsUK (N= 3,982) was too small to replicate the loci for such a heterogeneous phenotype as neck or shoulder pain. In particular, for the 4 SNPs chosen for replication, the power to achieve a *P* value < 0.05 in the given TwinsUK sample ranged between 11.6% to 15.8% (50).

The SNP heritability of neck or shoulder pain was 0.11. This is similar to that of back pain (0.11), greater than knee pain (0.08), and less than: hip pain (0.12), stomach or abdominal pain (0.14), headache (0.21), facial pain (0.24), and pain all over the body (0.31) (51). Further, the genetic correlation matrix among 8 pain phenotypes in the UK Biobank showed that the neck or shoulder pain and back pain shared the highest genetic correlation (rg = 0.83) (51).

The genetic correlation analysis results were perhaps to be expected. Like knee pain and back pain, which have been shown to be positively correlated with depression and neuroticism (51), neck or shoulder pain was correlated genetically and positively with some mental health and personality phenotypes. We also identified that neck or shoulder pain was genetically and negatively correlated with the age at which they have their first child, college completion, and years of schooling. This means that those who were older when they had their first child, those with more years of schooling, and those with completed college education were less likely to report neck or shoulder pain. These factors could be related to a number of factors including lifestyle, deprivation levels, and occupation. It is interesting to note that males are more likely to report neck or shoulder pain than females in the UK Biobank population. This is matched with the fact that males are more likely to have strenuous occupations. However, we should note, in general, female sex is a risk factor for neck or shoulder pain.

The primary limitation of this study is that different (albeit similar) case and control definitions were used in the discovery and replication cohorts. This was a consequence of the pre-determined phenotypic information that was present in the relevant cohorts. We defined neck or shoulder pain cases and controls based on the responses by UK Biobank participants to a specific pain question. This question focused on neck or shoulder pain occurrence during the previous month that was sufficient to cause interference with activity. The severity, frequency, and exact location of the neck or shoulder pain were not documented. Hence, our phenotyping should be considered as broadly defined. In the GS:SFHS and Twins UK cohorts, the disease status of participants was also self-reported, while cases were those who having neck or shoulder pain over the past 3 months, controls were those who could have neck or shoulder pain less than 3 months or have pain in other body sites. While in the UK Biobank, controls were defined as pain free for the past month. Differences between (albeit similar) the case and control definitions could have a negative impact on replication of the results while this impact is hard to evaluate Differences between the case and control definitions could have a negative impact on replication of the results although this impact would be hard to evaluate. There could also be some cases who report neck or shoulder pain as a result of underlying causes such as cancer and osteoarthritis in the neck and shoulder areas. Their impact to the results would be very limited due to the small numbers.

In summary, we have identified 3 loci of genome-wide significance (*P* < 5×10^−8^) associated with neck or shoulder pain in the UK Biobank dataset using a GWAS approach. Two of these loci were replicated weakly in the GS:SFHS cohort. Identification of these loci now provides a foundation for future work into understanding genetic roles and aetiology in neck or shoulder pain.

## ACKNOWLEDGEMENTS

We would like to thank all participants of the UK Biobank, Generation Scotland and TwinsUK cohorts who have provided necessary genetic and phenotypic information. This research has been conducted using the UK Biobank Resource under Application Number 4844.

Generation Scotland is grateful to all the families who took part, the general practitioners and the Scottish School of Primary Care for their help in recruiting them, and the whole Generation Scotland team, which includes interviewers, computer and laboratory technicians, clerical workers, research scientists, volunteers, managers, receptionists, healthcare assistants, and nurses.

## Data availability

The GWAS summary statistics of neck or shoulder pain can be accessed through https://figshare.com/articles/fourpainphenotype2/7699583

## Conflict of interest

The authors declare that they have no conflict of interest.

## Funding Sources

This study was mainly funded by the Wellcome Trust Strategic Award “Stratifying Resilience and Depression Longitudinally” (STRADL) with reference number 104036/Z/14/Z and by the GCRF academic exchange visits to China funded by the University of Dundee.

Generation Scotland: Scottish Family Health Studies (GS:SFHS) received core support from the Chief Scientist Office of the Scottish Government Health Directorates (CZD/16/6) and the Scottish Funding Council (HR03006).

TwinsUK is funded by the Wellcome Trust, Medical Research Council, European Union, Chronic Disease Research Foundation (CDRF), Zoe Global Ltd, the National Institute for Health Research (NIHR)-funded BioResource, Clinical Research Facility and Biomedical Research Centre based at Guy’s and St Thomas’ NHS Foundation Trust in partnership with King’s College London.

## Conflicts of Interest

None declared.

## AUTHOR CONTRIBUTIONS

WM organised the project, drafted the paper and contributed to the analysis. BWC and CH contributed to drafting the paper. MBF provided the TwinsUK replication. MA performed the main UK Biobank GWAS analysis. AC and CH contributed to GS:SFHS cohorts. HLH, HZ, XZ, CNAP, LC, TGH and FMKW provided essential comments. AM and BS organised the project and provided comments.

## Notes

https://figshare.com/articles/fourpainphenotype2/7699583

