## Supplementary figures and images for "A genome-wide association study finds genetic variants associated with neck or shoulder pain in UK Biobank"

### Supplementary Figure S1

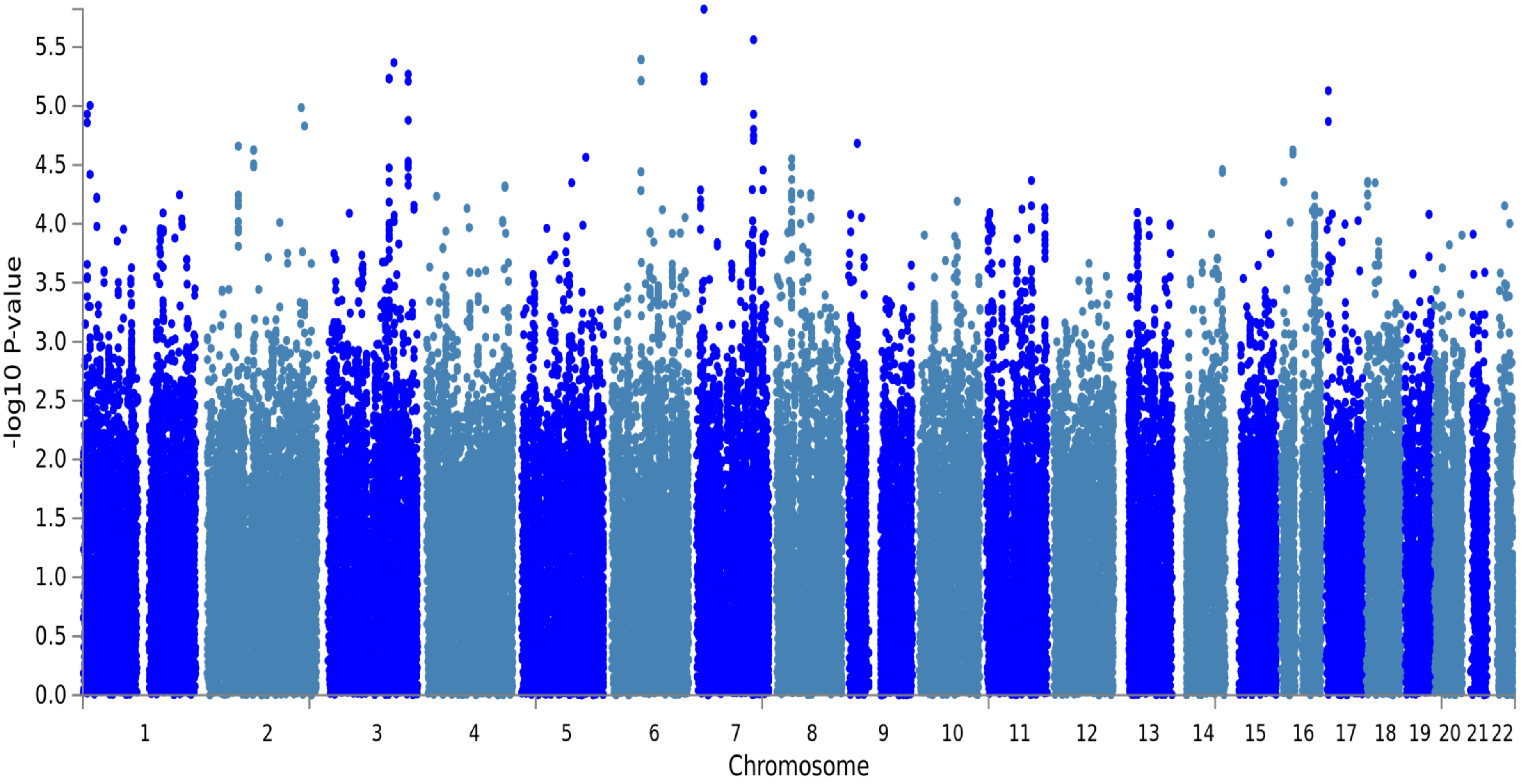


**Supplementary Figure S1:** the Manhattan plot of a GWAS on neck or shoulder pain using GS:SFHS (N=19,632)
