## Supplementary Table S1 for "A genome-wide association study finds genetic variants associated with neck or shoulder pain in UK Biobank"

| Gene | Chromosome | Start | Stop | NSNPS | NPARAM | ZSTAT | *P* |
|---|---|---|---|---|---|---|---|
| FOXP2 | 7 | 113726382 | 114333827 | 2311 | 132 | 6.6353 | 1.62E-11 |
| P2RY12 | 3 | 151055168 | 151102600 | 253 | 38 | 5.4674 | 2.28E-08 |
| PTBP1 | 19 | 797075 | 812327 | 121 | 36 | 5.4317 | 2.79E-08 |
| STEAP4 | 7 | 87905744 | 87936206 | 102 | 18 | 5.4316 | 2.79E-08 |
| MED12L | 3 | 150803484 | 151154860 | 1716 | 87 | 5.403 | 3.28E-08 |
| ZNF821 | 16 | 71893583 | 71929239 | 140 | 35 | 5.2693 | 6.85E-08 |
| LCOR | 10 | 98592017 | 98740800 | 499 | 62 | 5.2605 | 7.18E-08 |
| IST1 | 16 | 71879899 | 71962913 | 386 | 45 | 5.2235 | 8.78E-08 |
| TCTA | 3 | 49449639 | 49453908 | 13 | 7 | 5.2103 | 9.43E-08 |
| BSN | 3 | 49591922 | 49708978 | 346 | 40 | 5.1175 | 1.55E-07 |
| RHOA | 3 | 49396578 | 49450431 | 222 | 37 | 5.1144 | 1.57E-07 |
| DAG1 | 3 | 49506146 | 49573048 | 213 | 32 | 5.0012 | 2.85E-07 |
| NCAM1 | 11 | 112831997 | 113149158 | 1672 | 102 | 4.9828 | 3.13E-07 |
| ATXN1L | 16 | 71879894 | 71919171 | 152 | 38 | 4.9376 | 3.95E-07 |
| DCC | 18 | 49866542 | 51057784 | 7672 | 172 | 4.919 | 4.35E-07 |
| ILF3 | 19 | 10764937 | 10803093 | 141 | 32 | 4.9133 | 4.48E-07 |
| EYS | 6 | 64429876 | 66417118 | 11465 | 233 | 4.88 | 5.30E-07 |
| NICN1 | 3 | 49460379 | 49466759 | 11 | 5 | 4.8373 | 6.58E-07 |
| AGBL5 | 2 | 27265232 | 27293490 | 103 | 36 | 4.8085 | 7.60E-07 |
| GPX1 | 3 | 49394609 | 49396033 | 5 | 2 | 4.6874 | 1.38E-06 |
| SDCCAG8 | 1 | 243419320 | 243663394 | 854 | 84 | 4.6659 | 1.54E-06 |
| ZNF423 | 16 | 49521435 | 49891830 | 1794 | 187 | 4.6645 | 1.55E-06 |
| SLC44A2 | 19 | 10713133 | 10755235 | 217 | 37 | 4.653 | 1.64E-06 |
| CGREF1 | 2 | 27321757 | 27341995 | 84 | 26 | 4.6349 | 1.79E-06 |
| AP1G1 | 16 | 71762913 | 71843104 | 408 | 46 | 4.5696 | 2.44E-06 |
| TMOD2 | 15 | 52043758 | 52108565 | 375 | 27 | 4.5402 | 2.81E-06 |

NSNPS: Number of SNPs in the gene

NPARAM: Number of parameters used

**Supplementary Table S1.** The top 26 genes of the gene-based association analysis by MAGMA (integrated in FUMA)
